# Identification of a Δ11 desaturase from the arbuscular mycorrhizal fungus *Rhizophagus irregularis*

**DOI:** 10.1101/2020.01.13.903815

**Authors:** Henry Cheeld, Govindprasad Bhutada, Frederic Beaudoin, Peter J Eastmond

## Abstract

Arbuscular mycorrhizal fungi are oleaginous organisms and the most abundant fatty acyl moiety usually found in their lipids is palmitvaccenic acid (16:1^Δ11cis^). However, it is not known how this uncommon fatty acid species is made. Here we have cloned two homologs of Lepidopteran fatty acyl-CoenzymeA Δ11 desaturases from *Rhizophagus irregularis*. Both *DES1* and *DES2* are expressed in intraradicle mycelium and can complement the unsaturated fatty acid-requiring auxotrophic growth phenotype of the *Saccharomyces cerevisiae ole1Δ* mutant. DES1 expression leads almost exclusively to oleic acid (18:1^Δ9cis^) production, whereas DES2 expression results in the production of 16:1^Δ11cis^ and vaccenic acid (18:1^Δ11cis^). *DES2* therefore encodes a Δ11 desaturase that is likely to be responsible for the synthesis of 16:1^Δ11cis^ in *R. irregularis*.

## Introduction

Arbuscular mycorrhiza (AM) is the most common plant-microbe symbiotic association [1]. AM fungi are obligate biotrophs and receive organic carbon from their host plants in return for mineral nutrients [1]. Lipids are the major carbon currency in the AM fungal mycelium and they are transported to vesicles and spores where they are stored [2]. It was thought that AM fungi most likely synthesise their lipids *de novo* from sugars, which they receive from their host plant [3]. However, genomic analysis has suggested that AM fungi are fatty acid auxotrophs [4] and subsequent studies have shown that they rely on their host plant to supply them with long-chain fatty acyl moieties so that they can make fungal lipids [5-8]. The plant metabolic pathway that supplies fatty acyl moieties to AM fungi has been partially characterised, but it’s not yet clear precisely where this pathway ends and those of the fungus begin [5-8]. However, it is currently proposed that long-chain saturated fatty acyl moieties are most likely being transferred as 2-monoacylglycerols or free fatty acids [5-9].

The lipids in many (but not all) AM fungi are dominated by a single molecular species of monounsaturated fatty acid called 11-cis-palmitvaccenic acid (16:1^Δ11cis^), which can account for over 70 mol% of the fatty acyl moieties in their spores and is present mainly in the form of triacylglycerols [4,10-12]. 16:1^Δ11cis^ is unusual in that it contains a double bond at the ω5 (or Δ11) position and it has been used as a biomarker for arbuscular mycorrhization because it is not found in plants and it is rarely present in other soil microorganisms [10]. 16:1^Δ11cis^ has also been used in chemotaxonomy, because it is abundant in many AM fungi (Glomeromycota) but is lacking in certain species of the families Glomeraceae and Gigasporaceae [11].

It is thought that 16:1^Δ11cis^ is made in the intraradicle mycelium of AM fungi, but it is not known how [4, 12]. The discovery that AM fungi receive fatty acyl moieties from their host plant [5-8] also raises the possibility that 16:1^Δ11cis^ might be a product of plant metabolism. Understanding how and where 16:1^Δ11cis^ is made is therefore important to define how lipid metabolic pathways function within arbuscular mycorrhiza. Δ11 desaturases have previously been cloned from insects [13, 14] and marine diatoms [15], but we are not aware of any that have been characterised in fungi. The genomes of several AM fungi have now been sequenced, including *Rhizophagus irregularis* [16], which contains 16:1^Δ11cis^ [4]. A blastp search (https://www.ncbi.nlm.nih.gov/) of the *R. irregularis* genome using known Lepidopteran fatty acyl-CoenzymeA Δ11 desaturases [13, 14] revealed two potential homologues (DES1 and DES2). It is problematic to test the function of these genes in AM fungi because they are not amenable to genetic modification. We therefore characterized DES1 and DES2 by heterologous expression in *Saccharomyces cerevisiae* [13] and showed that *DES2* encodes a fungal Δ11 desaturase capable of synthesising 16:1^Δ11cis^.

## Materials and Methods

### Bioinformatic analysis of putative Δ11 desaturases

A blastp search was carried out in NCBI (https://www.ncbi.nlm.nih.gov/) on the *R. irregularis* DAOM197198 genome [16] using functionally-characterised Δ11 desaturase sequences from Lepidoptera [13, 14] and marine diatoms [15] as queries. All returned sequences with E scores <0.001 were compiled on a local server and aligned using Muscle v3.2 [17]. Two putative fatty acyl-CoA desaturases (GenBank accession numbers EXX76018 and EXX69612) were selected for further analysis and were named DES1 and DES2, respectively. The Kyte-Doolittle hydropathy scale with an amino acid window of 19 [18] and TMHMM v2.0 (http://www.cbs.dtu.dk/services/TMHMM/) [19] were used for hydropathy analysis and prediction of transmembrane helices (TMH). SignalP v4.0 (http://www.cbs.dtu.dk/services/SignalP) was used for identifying signal peptides at the N- and C-termini and for distinguishing these from TMH [20]. Searches for conserved domains within protein sequences were carried out using the NCBI conserved domain database (CDD) (https://www.ncbi.nlm.nih.gov/cdd) [21].

### Expression of DES1 and DES2 in *S. cerevisiae*

The open reading frames of *DES1* and *DES2* were codon optimised for expression in *S. cerevisiae* by Genscript, synthesised and supplied in the pUC-57 vector. *DES1* and *DES2* were then excised using *BamHI* and *SalI* restriction sites and ligated into pHEY1 [22], for expression under the constitutive *TEF1* promoter. pHEY-DES1, pHEY-DES2 and pHEY-EVC (empty vector) were transformed into *S. cerevisiae* [23] wild type strain DTY-11a (*MATa, leu2-3, leu2-12, trp1–1, can1–100, ura3–1, ade2–1*) and *ole1Δ* knockout strain AMY-3α (*MATα, ole1Δ::LEU2, trp1–1, can1–100, ura3–1, ade2–1*) [24] and colonies were selected on synthetic Dexterose (SD) minimal medium agar plates lacking uracil. The SD minimal medium used for selection of yeast transformants and culture cultivations consisted of 6.9 g L^-1^ yeast nitrogen base without amino acids (Formedium), 1.92 g L^-1^ yeast synthetic drop-out medium supplements minus uracil (Merck), 40 mg L^-1^ of adenine (Merck) and 20 g L^-1^ glucose (Merck) as sole carbon source. 10 mL cultures were grown overnight in SD minimal media to optical density (OD) of 0.5 to 1 at 600 nm, then used to inoculate in 100 mL of SD minimal media to a starting OD600 of 0.1. The 100 mL cultures were then incubated at 30 °C and shaken at 200 rpm for 72h. *ole1Δ* cultures were supplemented with 1mM odd or even chain monounsaturated fatty acids (MUFAs), emulsified in 1% (v/v) tergitol (Merck). *ole1Δ* was also grown on SD minimal media agar plates containing fatty acids.

### Lipid extraction and analysis

Cultures were normalised for cell volume based on OD600 measurements and the cells were pelleted by centrifugation at 4000 rpm, the supernatant discarded, and the pellet frozen in liquid nitrogen and stored at -80°C. Fatty acid methyl esters (FAMEs) were prepared from the cell pellets by transmethylation in 1 ml of methanol/toluene/dimethoxypropane/H_2_SO_4_ (66:28:2:1 by volume) at 80°C for 40 min. 0.5 ml hexane and 1 ml KCl (0.88% w/v) were added and the contents vortexed, and centrifuged, before transferring the upper hexane phase to a fresh vial. Extraction with hexane was repeated twice to ensure extraction of all FAMEs and the three extracts pooled. The FAMEs were dried down under N_2_ and reconstituted in 0.5 ml heptane, and 75 µL taken for analysis by gas chromatography (GC) coupled to mass spectrometry (MS) or flame ionisation detector (FID). Position of double bonds in monounsaturated FAMEs were determined by preparing dimethyl disulphide (DMDS) adducts [25]. 100-1000 ng FAMEs (in 50 µL hexane), 5 µL 50 mg ml-1 iodine in diethyl ether, and 50 µL DMDS, were combined and vortexed followed by heating at 40°C for 15 hrs. 5 µL 5% (w/v) sodium thiosulphate and 200 µL hexane were added, vortexed and centrifuged to separate the phases. The hexane layer was removed, dried under N_2_, reconstituted in 50 µL heptane for analysis by GC-MS. Separation of FAMEs and DMDS adducts was performed by 6890N Network GC System (Agilent) fitted with a 30 m × 0.25 mm, 0.25 µm film thickness, HP1-MS-UI capillary column (Agilent). 1 µL FAME/DMDS adducts were injected (splitless) at 280°C and He used as the carrier gas (0.6 mL min^-1^) at a constant flow. The oven program was as follows: 70°C (1 min), 40°C min^-1^ ramp to 150°C, 4°C min^-1^ ramp to 300°C (2 mins), 325°C (18 mins). For FAME/DMDS adduct identification, GC was coupled to a 5975B mass selective detector (Agilent) with a 3.5 min solvent delay, on constant scan mode 42 – 500 m/z. The detection and quantification of FAMEs by GC-FID was also performed, using a DB-23 capillary column (Agilent) as described previously [26].

## Results

### Identification of putative Δ11 desaturases

To identify candidate Δ11 desaturases from AM fungi we performed a blastp search of the *R. irregularis* DAOM197198 genome [16] using characterised insect [13, 14] and marine diatom [15] protein sequences. Two genes designated *DES1* and *DES2* were identified that encode proteins that share substantial (>35%) sequence identity with the archetypal palmitoyl (16:0)-CoA Δ11 desaturase from *Trichoplusia ni* (GenBank accession number AAD03775) [13]. Comparison of the amino acid sequences (Fig. 1) revealed that DES1 and DES2 contain a membrane desaturase-like conserved domain (cl00615) [21] which includes three His-box motifs, characteristic of desaturases and essential for their catalytic activity [27]. In addition, DES1 and DES2 also contain a cytochrome b5-like heme-binding conserved domain (cl34968) [21] at their C-terminus, featuring a HPGG motif (Fig. 1) that is characteristic of the fusion between deasturases and cytochrome b5 [28]. Both insect and mammalian fatty acyl-CoA desaturases lack cytochrome b5-like C-terminal extensions, but they are present in fungal counterparts such as the *S. cerevisiae* Δ9 desaturase Ole1p [24, 29]. Transmembrane helices (TMHs) predicted by the Kyte-Doolittle hydropathy scale [18] and TMHMM [19] were in qualitative agreement, and both algorithms placed the N- and C-termini of DES1 and DES2 in the cytosol, consistent with the topology of mammalian stearoyl (18:0)-CoA desaturases (SCDs) [30]. Four TMHs were identified, which are typical features of membrane-bound desaturases [30] and are consistent with the crystallographic structures of SCDs [31, 32].

**Figure 1.**
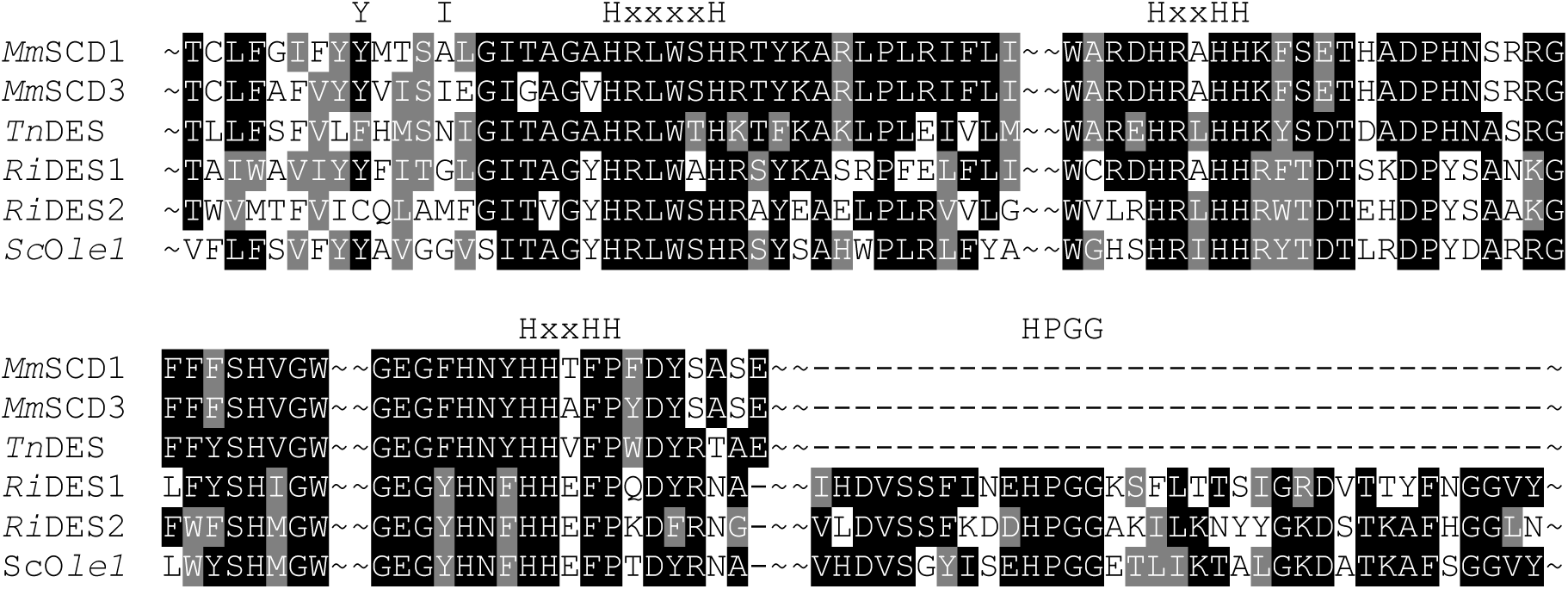
Conserved regions of *R. irregularis* (*Ri*) DES1 and DES2 aligned with *Trichoplusia ni* (*Tn*) Δ11 desaturase and other functionally-characterised fatty acyl-CoA desaturases from *M. musculus* (*Mm*) and *S. cerevisiae* (*Sc*). The three conserved His-boxes [H(x(n)(H)H] of the desaturase domain and the HPGG box in the C-terminal cytochrome b5-like fusion domain are highlighted. Residues Tyr 133 and Ile 137 that face the substrate binding pocket in *Mm*SCD3 are also highlighted.

### Expression of *DES1* and *DES2* in *R. irregularis*

To investigate whether *DES1* and *DES2* are expressed in *R. irregularis* we analysed a RNA-sequencing data set that includes structures from both asymbiotic and symbiotic stages of the AM fungal life cycle such as germ tubes, runner hyphae, intraradical mycelium, arbuscules, branched absorbing structures, immature and mature spores [33]. A search for the corresponding transcripts of *DES1* and *DES2* within this data set revealed that both genes are expressed in all seven AM fungal structures, but *DES2* appears to be the more strongly expressed of the two genes, particularly in intraradical mycelium, arbuscules and spores (Table 1). A desaturase responsible for producing 16:1^Δ11cis^ in *R. irregularis* should be expressed in these structures since this fatty acyl moiety is most abundant in triacylglycerol that accumulates first in lipid droplets that form in the intraradical mycelium proximal to arbuscules [2, 34].

**Table 1.** Transcript abundance of *DES1* and *DES2* in different structures of *R. irregularis*.

| Structure | GT | RH | IRM | ARB | BAS | IS | MS |
| --- | --- | --- | --- | --- | --- | --- | --- |
| *DES1* | 3.04 | 6.51 | 7.99 | 10.93 | 9.30 | 11.68 | 13.71 |
| *DES2* | 5.88 | 7.66 | 6.80 | 7.90 | 6.80 | 8.22 | 7.22 |

RNA-sequencing data are derived from Kameoka et al., [33] and are expressed as mean Log2 FPKM (fragments per kilobase of exon per million reads mapped). GT, germ tubes; RH, runner hyphae; IRM, intraradical mycelium; ARB, arbuscules; BAS, branched absorbing structures; IS, immature spores; MS, mature spores.

### Functional analysis of DES1 and DES2 by expression in *S. cerevisiae*

To test the enzymatic function of DES1 and DES2 we transformed wild type *S. cerevisiae* and the desaturation-deficient *ole1Δ* knock-out strain [24, 29] with the high copy number plasmids pHEY-DES1 and pHEY-DES2, designed to express the two genes under the control of the strong constitutive translational elongation factor EF-1α (*TEF1*) promoter [22]. The *ole1Δ* strain is completely deficient in fatty acid desaturation and can only grow on media that is supplemented with exogenous long-chain unsaturated fatty acids [24, 29]. A plate test of *ole1Δ* harbouring either pHEY-DES1 or pHEY-DES2 showed that cell growth could be rescued by expression of DES1 or DES2 (Fig. 2), suggesting that both proteins can function as desaturases [24, 29].

**Figure 2.**
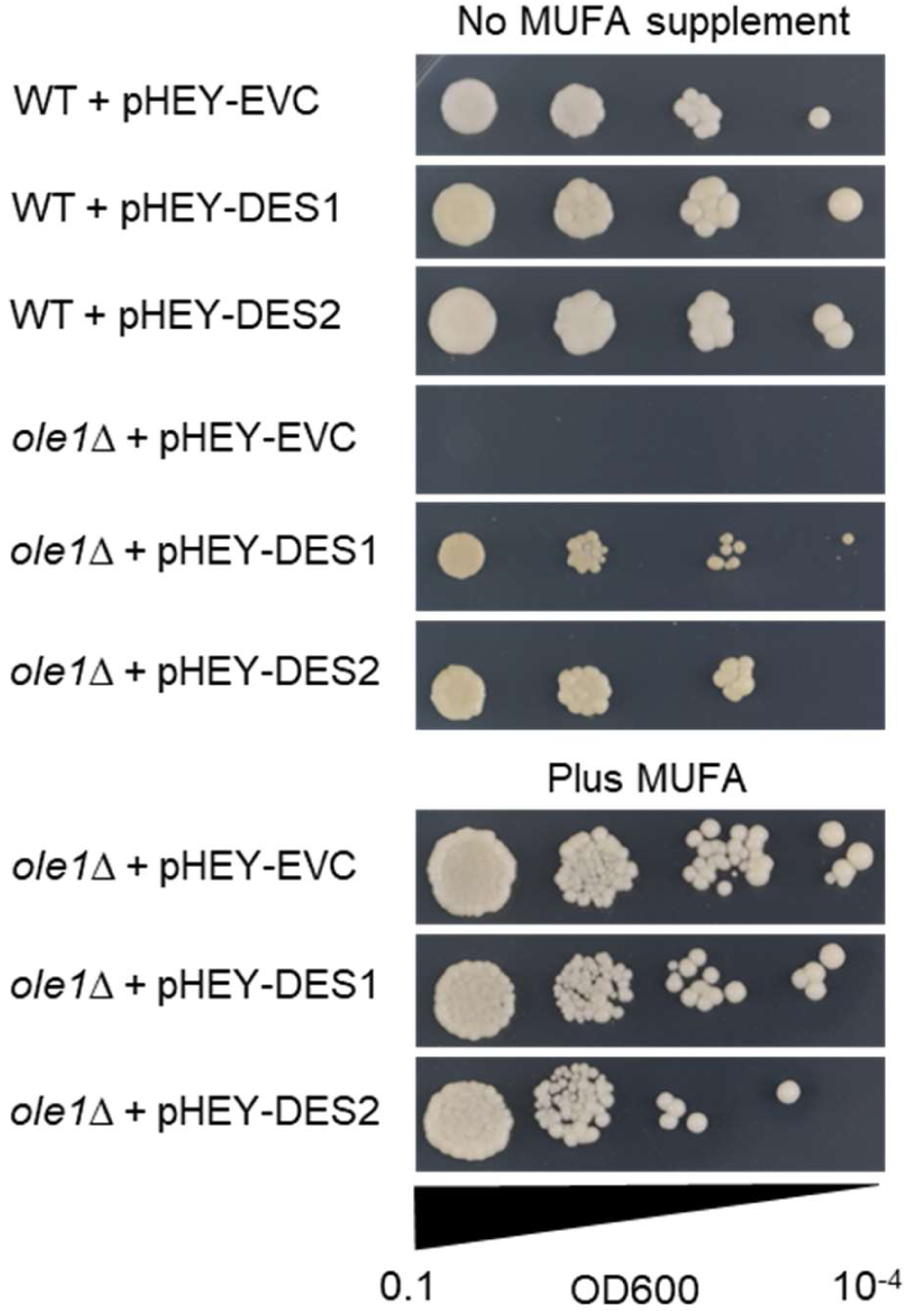
Plate test illustrating the ability of DES1 and DES2 to rescue the unsaturated fatty acid auxotrophic phenotype of *S. cerevisiae ole1Δ*. A 0.1 OD600 culture was successively diluted 10-fold to 10^−4^ and 2 µL drops added to plates with or without a monounsaturated fatty acid (MUFA) supplement, using 1 mM C15:1^Δ10cis^. Image was taken after 72 hrs growth at 30°C.

Analysis of the fatty acyl composition of lipids from wild type *S. cerevisiae* cells expressing DES1 revealed that there was no change in the molecular species that were produced (Fig. 3). However, there was a significant (P > 0.05) increase in the relative abundance of oleic acid (C18:1^Δ9cis^), as compared to the empty vector control (EVC) (Fig. 3; Table S1). By contrast, DES2 expression in wild type cells led to the appearance of two major new molecular species of fatty acyl moiety (Fig. 3), which GC-MS analysis indicated were isomers of 16:1 (m/z 268) and 18:1 (m/z 296). Further analysis of the double bond positions by extraction of the molecular ions of dimethyl disulphide (DMDS) adducts [25] revealed the characteristic fragment ions of 16:1^Δ11cis^ (m/z 117, 245) and 18:1^Δ11cis^ (m/z 144, 245) (Fig. S1). Small amounts of 13-*cis*-octadeccenoic (18:1^Δ13cis^) (m/z 117, 273) were also detected (Fig. 3; Table S1; Fig. S1). Further analysis of the fatty acyl composition of *ole1Δ* cells expressing DES1 or DES2 confirmed that, with the substrates that are available, DES1 preferentially produces C18:1^Δ9cis^ over C16:1^Δ9cis^, whereas DES2 produces 16:1^Δ11cis^ and to a lesser extent 18:1^Δ11cis^ (Fig. 3; Table S1).

**Figure 3.**
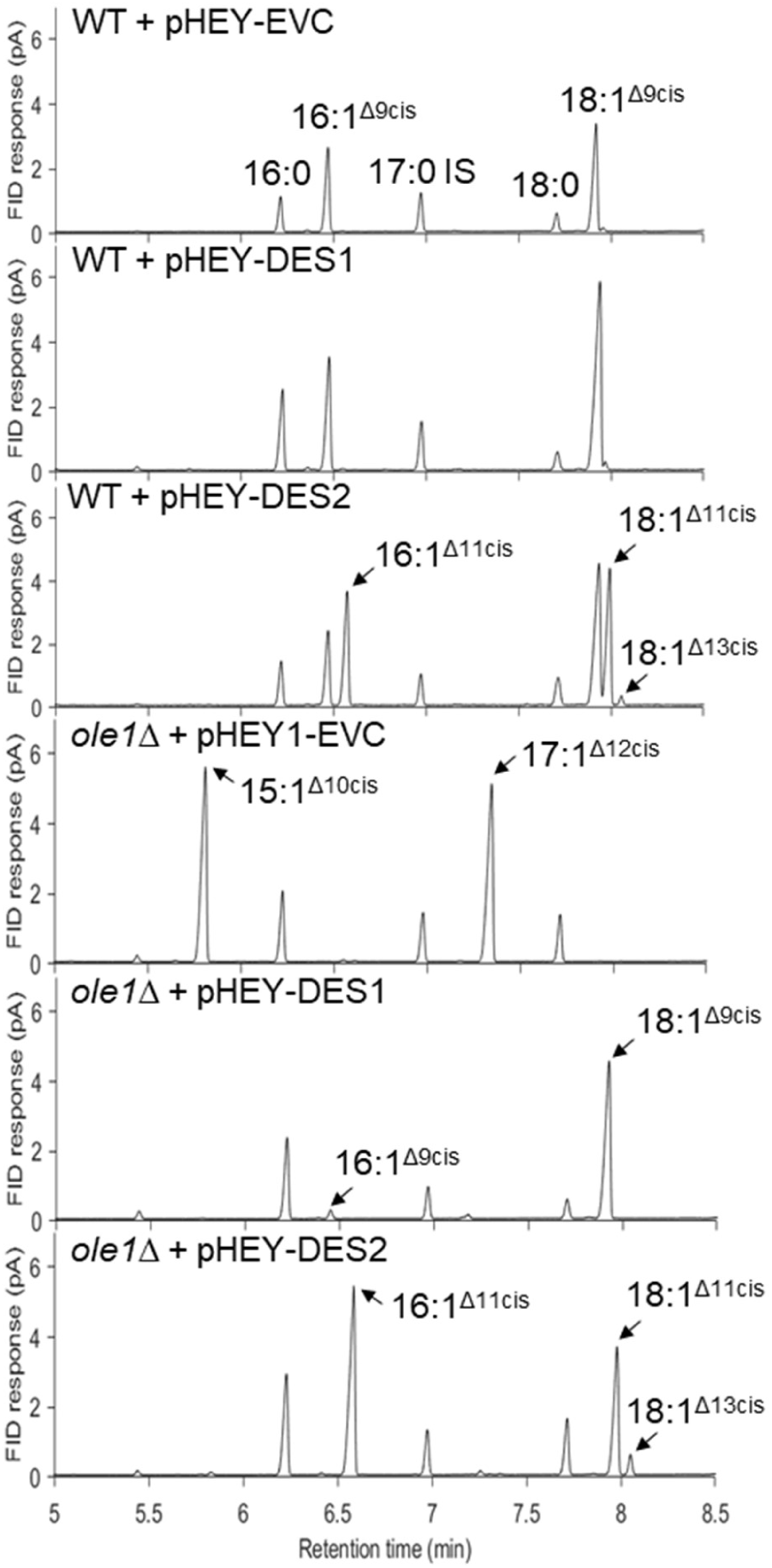
GC-FID analysis of FAMEs derived from lipid extracts of wild type (WT) or *ole1Δ* cultures harbouring pHEY vectors, either as EVC or containing DES1 or DES2. For *ole1Δ* + pHEY1-EVC an odd chain MUFA supplement (15:1^Δ10cis^) was used to complement the *ole1Δ* phenotype and the 17:1^Δ12cis^ is an elongation product of 15:1^Δ10cis^. 17:0 was included as an internal standard (IS). The individual GC-FID traces are representative of three replicates.

In WT *S. cerevisiae* cells, trace amounts of 16:1^Δ11cis^ and 18:1^Δ11cis^ were also detected (Table S1). 16:1^Δ11cis^ is known to be a product of 9-cis-myristoleic acid (14:1^Δ9cis^) elongation by Elo1p [35] and 18:1^Δ11cis^ is most likely an elongation product of 16:1^Δ9cis^. 16:1^Δ11cis^ elongation is also likely to explain the small amounts of 18:1^Δ13cis^ detected in both WT and *ole1Δ* cells expressing DES2. To test this hypothesis *ole1Δ* cells expressing DES2 were supplemented with 16:0 or 18:0 free fatty acids to increase the respective amounts of substrate available for desaturation. The addition of 16:0 resulted in a significant increase in 16:1^Δ11cis^ and 18:1^Δ13cis^ (P > 0.05), which is consistent with a precursor-product relationship (Table S1). Addition of 18:0 resulted in a significant increase in 18:1^Δ11cis^ (P > 0.05), but not in 18:1^Δ13cis^ (Table S1), suggesting that these monounsaturated fatty acids (MUFAs) are not products of the same substrate. Taken together, these data suggest that the 18:1^Δ13cis^ is not a direct product of 18:0 desaturation, but of 18:1^Δ11cis^ elongation.

## Discussion

Our data show that *R. irregularis* contains two desaturases that share sequence similarity with fatty acyl-CoA Δ11 desaturases from Lepidoptera [13], but also possess a cytochrome b5-like C-terminal extension characteristic of their fungal Δ9 counterparts [24, 29]. DES1 and DES2 are both expressed in the intraradicle mycelium where 16:1^Δ11cis^ is thought to be synthesised [4, 12]. DES1 and DES2 also both function as desaturases, since they can complement the MUFA-deficient phenotype of the *S. cerevisiae ole1Δ* mutant [13, 29]. However, analysis of their products shows that DES1 synthesises 18:1^Δ9cis^ and a little 16:1^Δ9cis^, whereas DES2 synthesises 16:1^Δ11cis^ and 18:1^Δ11cis^. DES2 activity is therefore most likely to account for the high levels of 16:1^Δ11cis^ that accumulate in *R. irregularis* [4, 11, 12].

Desaturases are classified based on their ability to recognise either the ω (methyl) or Δ (carboxyl) end of the fatty acyl moiety for insertion of the double bond [36]. The ability of DES1 and DES2 to produce Δ9 and Δ11 fatty acids using substrates with different chain lengths (C16 and C18) suggests that both are front-end desaturases that count carbon atoms from the carboxyl-terminus for insertion of the double bond. The structural basis of chain length specificity has been studied previously in fatty acyl-CoA desaturases [31]. The substrate binding channel of *Mus musculus* SCD1 is capped by Tyr 133, which is located on the second transmembrane helix and blocks access of acyl chains longer than C18 [31, 37]. DES1 also possess Tyr in the corresponding position (Fig. 1). One helical twist above Tyr 133 in *M. musculus* SCD3, and therefore facing the binding pocket, is Ile 137 [31]. Mutant analysis suggests that when Ile 137 is substituted for Ala (as is found in SCD1) SCD3 substrate preference changes from C16 to C18 [31]. Ile has a bulkier side chain than Ala and may therefore shorten the substrate channel [31]. DES1 has Gly in this position (Fig. 1), which has a small side chain. DES2 has Met in this position (Fig. 1), which has a slightly larger side chain. The residues occupying these positions might therefore explain why DES1 prefers a C18 substrate and DES2 prefers C16.

Although 16:1^Δ11cis^ is highly abundant in *R. irregularis*, levels of 18:1^Δ11cis^ are much lower [4,12]. Given that DES2 can synthesise both MUFAs in *S. cerevisiae*, it is possible that the high 16:1^Δ11cis^ to 18:1^Δ11cis^ ratio found in *R. irregularis* could be a result of a relative difference in substrate availability. It is thought that *R. irregularis* receives fatty acyl moieties from its host plant that are mainly C16, or shorter [5-9] and so these substrates might be more abundant. *R. irregularis* also contains a comparatively low level of 18:1^Δ9cis^ [4, 12] that is likely to be produced by DES1, given its activity in *S. cerevisiae*. In addition to *R. irregularis*, 16:1^Δ11cis^ is present in many Glomeromycota and putative orthologues of DES2 can also be found in the *R. diaphanous, R. clarus, R. cerebriforme* and *Gigaspora rosea* genomes [38], but not in those of non-mycorrhizal fungi. Interestingly, *G. rosea* is one of the species from the family Gigasporaceae that does not contain 16:1^Δ11cis^ [11, 12]. It is therefore possible that *G. rosea* DES2 either has a different activity (i.e. is not a Δ11 desaturase) or is not expressed. At present it is not known why many Glomeromycota make 16:1^Δ11cis^ and some do not. The identification of DES1 and DES2 may help in future studies to better understand the physiological role of the different molecular species of MUFAs found in AM fungi.

## Abbreviations

AM: arbuscular mycorrhizal
CoA: CoenzymeA
16:1^Δ11cis^: 11-cis-palmitvaccenic acid
18:1^Δ11cis^: 11-cis-vaccenic acid
SCD: stearoyl-CoA desaturase
TEF1: translational elongation factor EF-1α

## Acknowledgements

We thank Charles Martin for providing the *ole1Δ* mutant. This work was funded by the UK Biotechnology and Biological Sciences Research Council through grant BB/P012663/1.

## Author contributions

P.J.E. conceived the research; H.C., G.B. and F.B. performed the research; H.C., G.B. and F.B. analysed data; H.C. and P.J.E. wrote the paper.

