## Supplemental data for "Identification of a Δ11 desaturase from the arbuscular mycorrhizal fungus *Rhizophagus irregularis*"

Supporting information

Table S1. Fatty acid composition of WT or *ole1Δ* cultures expressing DES1 or DES2.

| Fatty acid | 16:0 | $16:1^{\Delta 9cis}$ | $16:1^{\Delta 11cis}$ | 18:0 | $18:1^{\Delta 9cis}$ | $18:1^{\Delta 11cis}$ | $18:1^{\Delta 13cis}$ |
|---|---|---|---|---|---|---|---|
| WT + pHEY-EVC | 12.15 ±0.01 | 33.61 ±0.02 | 0.22 ±<0.01 | 9.27 ±0.02 | 42.64 ±0.03[a] | 2.11 ±0.01 | nd |
| WT + pHEY-DES1 | 16.72 ±0.01 | 22.99 ±0.01 | 0.23 ±<0.01 | 3.70 ±0.01 | 54.01 ±0.03[a] | 2.34 ±0.01 | nd |
| WT + pHEY-DES2 | 7.70 ±0.01 | 15.15 ±0.02 | 20.90 ±<0.01 | 5.29 ±<0.01 | 30.03 ±<0.01 | 19.86 ±0.03 | 1.04 ±<0.01 |
| *ole1Δ* + pHEY-EVC (plus 1mM $15:0^{\Delta 10cis}$) | 62.80 ±<0.01 | nd | nd | 37.20 ±<0.01 | nd | nd | nd |
| *ole1Δ* + pHEY-DES1 | 28.29 ±0.01 | 2.64 ±<0.01 | nd | 6.13 ±<0.01 | 62.94 ±0.01 | nd | nd |
| *ole1Δ* + pHEY-DES2 | 22.29 ±0.01 | nd | 35.21 ±0.02[b] | 10.64 ±<0.01 | nd | 28.64 ±0.01[d] | 3.31 ±<0.01[c] |
| *ole1Δ* + pHEY-DES2 (plus 1mM 16:0) | 27.66 ±0.03 | nd | 53.61 ±0.01[b] | 4.12 ±0.01 | 0.14 ±<0.01 | 8.99 ±0.01 | 5.45 ±<0.01[c] |
| *ole1Δ* + pHEY-DES2 (plus 1mM 18:0) | 12.78 ±0.01 | nd | 30.16 ±0.03 | 15.24 ±0.01 | nd | 40.42 ±0.06[d] | 1.33 ±<0.01 |

FAMEs were quantified using GC-FID analysis and their identities were determined by GC-MS analysis of DMDS adducts. nd is not detected. Values are expressed as a percentage of the total and are the mean ±SE of measurements made on cells from three separate cultures. a, b, c and d denote specific pairs of values discussed in the text that are significantly different (P > 0.05; two-tailed Student's t test).

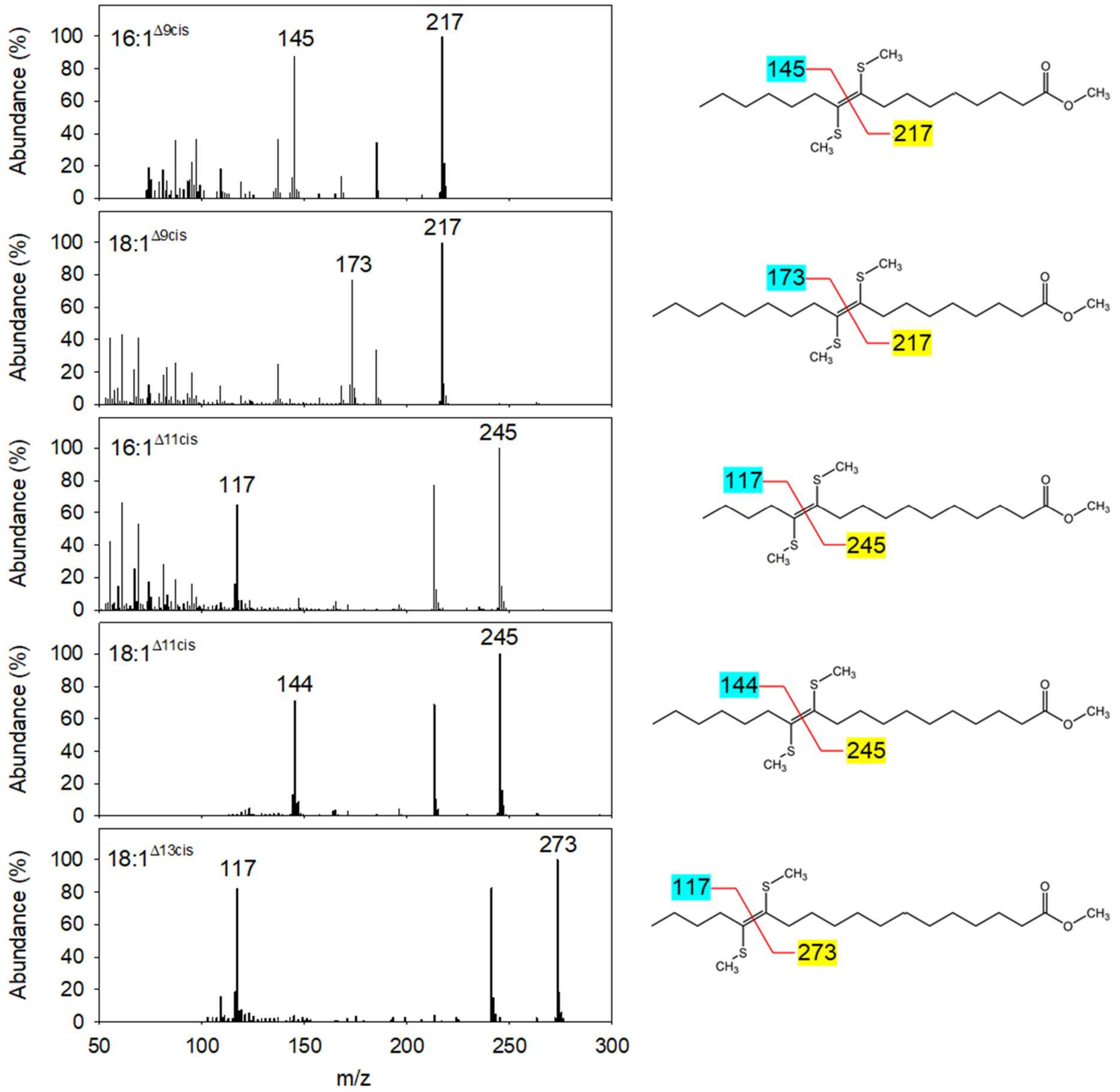


Figure S1. Mass spectra of DMDS FAME adducts. Annotated ions represent diagnostic fragment ions of DMDS adducts used for locating the double bond. Examples of 16:1$^{\Delta 9cis}$ and 18:1$^{\Delta 9cis}$ from *ole1Δ* + pHEY-DES1 and 16:1$^{\Delta 11cis}$, 18:1$^{\Delta 11cis}$ and 18:1$^{\Delta 13cis}$ from *ole1Δ* + pHEY-DES2.
